# Metabolic cell death of labile iron deficiency as a vulnerability of head and neck squamous cell carcinoma evaded by BACH1

**DOI:** 10.1101/2025.04.07.647470

**Authors:** Masahiro Rokugo, Kazuki Nakamura, Mitsuyo Matsumoto, Akari Endo, Long Chi Nguyen, Hironari Nishizawa, Ryohei Shima, Hitomi Kashiwagi, Akira Okoshi, Ayako Nakanome, Ryo Funayama, Keiko Nakayama, Takenori Ogawa, Takaaki Abe, Yukio Katori, Kazuhiko Igarashi

**Author notes:** Corresponding author (KI,; MM,).

## Abstract

Head and neck squamous cell carcinoma (HNSCC) remains difficult to treat due to the lack of molecularly targeted drugs with high anti-tumor efficacy and safety. To investigate the involvement of the transcription factor BACH1, which is known to promote the metastasis of various cancers, in the malignant properties of HNSCC cells, we examined the effects of BACH1 depletion with short interfering RNAs in human HNSCC cell lines. We found that knockdown of BACH1 induced cell death due to depletion of intracellular labile iron which is not tightly bound to protein. Mitochodrial electron transfer chain activity was severely reduced but efficiently resuced with supplementation of iron in the medium. By combining chromatin immunoprecipitation-sequence and RNA-sequence analyses, we found that BACH1 represses ferritin genes in HNSCC cells. BACH1 knockdown was found to enhance the effect of tipifarnib, a selective inhibitor of farnesyltransferase required for HRAS activation, resulting in efficient inhibition of proliferation of HNSCC cell lines in vitro. These results indicate that BACH1 is essential for maintaining the amount of labile iron in HNSCC cells to evade metabolic cell death of iron deficiency and to proliferate. BACH1 can be a target for conferring Tipifarnib sensitivity to HNSCC cells.

## Introduction

Head and neck cancer was the eighth most common cancer worldwide (745,000 new cases and 365,000 deaths in 2020) (1). More than 90% of head and neck cancers are squamous cell carcinomas (HNSCC). The prognosis for HNSCC patients has not improved; 50-55% cases die from the disease within 5 years and the treatment effect seems to have reached a plateau (2). While multidisciplinary treatment combining surgical and non-surgical treatments is important, for many years, a combination of platinum-based chemotherapy has represented the only available option for first-line therapy. Cetuximab, a human monoclonal antibody against the epidermal growth factor receptor (EGFR), is the only clinically available molecularly targeted drug for HNSCC.

However, it is not superior to standard treatment in terms of its antitumor effect and adverse events, and the indications for Cetuximab in clinical practice are very limited (3, 4). Unfortunately, the only available treatments for recurrent/metastatic HNSCC are medications and palliative radiation therapy and/or surgery for the purpose of prolonging life or maintaining quality of life. In recent years, immune checkpoint inhibitors targeting PD-1 has been introduced for recurrent/metastatic HNSCC, but the benefits are limited to only a fraction of patients (5, 6). Recently, Tipifarnib, a specific inhibitor of farnesyltransferase of RAS proteins, has been tested in the clinical trial of HNSCC with a mutation of the *HRAS* gene (7–9). Tipifarnib is particularly effective in HNSSCC with an *HRAS* variant allele frequency (VAF) of 20% or higher (9). However, because a substantial fraction of tumors are with VAF less than this value, one of the questions is to find a way to augment the effectiveness of Tipifarnib. To advance the treatment of HNSCC, detailed understanding of molecular mechanisms critical for the survival and proliferation of the cancer cells is needed.

The transcriptional factor BTB and CNC homolog 1 (BACH1) has been reported to contribute to the malignant progression of various carcinomas such as melanoma, colorectal cancer (10), lung cancer (11), pancreatic cancer (12), and esophageal cancer (13). It has also been suggested that BACH1 may be a therapeutic target in cancer (14, 15). Considering that transformation and tumor formation of mouse embryonic fibroblasts (MEFs) with active mutant *H*r*as* is dependent on *Bach1* (16), BACH1 may facilitate malignant progression of cancers with *RAS* mutations, including HNSCC.

One of the biochemical functions regulated by BACH1 is iron metabolism. Among metallic elements, iron is essential for living organisms as a constituent of hemoglobin for the transport of oxygen, diverse proteins in energy metabolism including electron transfer chain in the mitochondria, and enzymes including oxidoreductases and those for DNA synthesis (17). BACH1 represses the expression of heme oxygenase-1 (HO-1 encoded by *HMOX1*), which mitigates oxidative stress and promotes recycling of iron by degrading heme. BACH1 also represses ferritin genes (*FTH1*, *FTL*) (18–22) and genes in the glutathione synthesis pathway (23), promoting ferroptosis, an iron-dependent cell death. Iron homeostasis is attracting attention as a target of cancer therapy (24–31), including the use of iron chelators in combination with existing chemotherapeutic agents to enhance the antitumor effect (32, 33). While BACH1 is involved in cellular iron homeostasis and is involved in malignant features of diverse types of cancer cells as described above, it is not clear whether these two roles are mechanistically connected with each other. In addition, the function of BACH1 in HNSCC has not been reported yet. In this study, we tried to investigate the function of BACH1 in HNSCC with a focus on iron. We further investigated whether BACH1 is a target to increase the efficacy of tipifarnib for HNSCC.

## Material and method

### Cell culture and reagent

In this study, two human head and neck cancer cell lines (HSC-2 and Ca9-22) were used as patient-derived cell lines for oral cancer squamous cell carcinoma cell lines. Both cell lines were obtained from the Medical Cell Resource Center affiliated with the Institute of Development, Aging and Cancer, Tohoku University. 10% fetal bovine serum (Fetal Bovine Serum, FBS, Sigma Aldrich), 100 U/ml penicillin (GIBCO, NY, USA) and 100 µg/ml streptomycin (GIBCO) in a medium supplemented with 10 mM HEPES (GIBCO), and the culture was maintained under 37L, 5% COL conditions. For HSC-2 with BACH1 overexpression system using TET-on system, fetal bovine serum was replaced with Tet-System Approved FBS (Thermo Fisher Scientific Inc., MA, USA) and used at 10% of medium volume.

### RNA interference

BACH1 siRNA (Stealth RNAi™ siRNA Duplex Oligoribonucleotides, Invitrogen.) and Stealth RNAi™ siRNA Negative Control, Middle GC (Thermo Fisher Scientific Inc.) were used as controls. Lipofectamine RNAiMAX (Thermo Fisher Scientific Inc.) was used to introduce siRNA by the lipofection method according to the product manual.

The sequence of the siRNA used is as follows.

siBACH1-1; 5’-GGUCAAAGGACUUUCACAACAUUAA-3’
siBACH1-3; 5’-GCAUUUGGAACUGACAGAGUCCGUA-3’

### RNA extraction and cDNA synthesis

RNA extraction was performed using the RNeasy Plus Mini Kit (Qiagen, Germany) according to the product instructions. The concentration of the extracted RNA was measured using an absorbance meter (Thermo Fisher Scientific Nano Drop 1000 Spectrophotometer). Reverse transcription reactions were then performed from 350 to 1,000 ng of extracted RNA using SuperScript III Reverse Transcriptase® (Thermo Fisher Scientific) and Random Primers (Thermo Fisher Scientific).

### Quantitative real time PCR (qPCR)

Quantified real time PCR was performed by the intercalator method using SYBR GreenL. LightCycler Fast Start DNA Master SYBR Green I (Roche, Switzerland) and performed according to the product instructions. Each gene-specific primer and the cDNA of each sample were mixed and measured on a quantitative PCR machine LightCycler®96 (Roche). The gene of ribosomal Protein L13a (*RPL13A*) was used as the internal standard gene. The sequence of each primer is as follows.

#### BACH1

(Forward) 5’-AATCGTAGGCCAGGCTGATG-3’

(Reverse) 5’-AGCAGTGTAGGCAAACTGAA-3’

#### RPL13A

(Forward)5’-TCGTACGCTGTGAAGGCATC-3’

(Reverse) 5’-GTGGGGCAGCATACCTCG-3’

#### FTL

(Forward) 5’-TACGAGCGTCTCCTGAAGATGC-3’

(Reverse) 5’-GGTTCAGCTTTTTCTCCAGGGC-3’

#### FTH1

(Forward) 5’-TGAAGCTGCAGAACCAACGAGG-3’

(Reverse) 5’-GCACACTCCATTGCATTCAGCC-3’

#### TFRC

(Forward) 5’- ATCGGTTGGTGCCACTGAATGG-3’

(Reverse) 5’- ACAACAGTGGGCTGGCAGAAAC-3’

### Cell counting assay

For cell proliferation analysis after RNA interference, cells (3 × 10L cells per well) were seeded in 3.5 cm culture dishes and transfected with 30 pmol of siRNA using Lipofectamine RNAimax the next day. The number of adherent cells was measured at 24, 48, and 72 hours after transfection. After collected cells using trypsin, and the number of living cells was manually measured under a microscope.

### Cell viability assay

The WST method was used for cell viability assay. Cell Counting Kit-8 (Dojindo Labolatories, Japan) was administered at 7µl per sample. One hour after Cell Counting Kit-8 administration, absorbance (450 nm) was measured using iMark™ Microplate Reader (Bio-rad).

### Western blotting

Total protein extracts of heat-denatured cells were used as samples, which were applied to 5.0%-20% precast gels for electrophoresis (Longevity Gel®, Oriental Instruments), After SDS-PAGE, the samples were transferred to polyvinylidene fluoride (PVDF) membranes (Millipore IPVH00010). The transferred PVDF membrane was immersed in 4% skim milk (Wako Pure Chemical Co., Osaka, Japan) diluted with T-TBS (TBS (25 mM Tris-HCl (pH 7.4), 137 mM NaCl, 3 mM KCl) containing 0.05% Tween 20) and blocked. Primary antibodies were reacted with the above blocking solution or bovine serum albumin (BSA) solution (TBS solution containing 5% BSA and 0.1% Tween 20) by diluting various antibodies 1000-fold (except GAPDH) or 5000-fold (GAPDH only), followed by shaking at 4°C overnight. Using either Rabbit IgG HRP Linked Whole Ab (GE healthcare NA934) or Mouse IgG HRP Linked Whole Ab (GE healthcare NA931) as a secondary antibody, dilute the antibody 5000-fold (BACH1 and GAPDH) or 1000-fold (other than BACH1 and GAPDH) in T-TBS containing 4% skim milk or BSA solution. The antibody was reacted with Supersignal West Pico Plus Chemiluminescent Substrate (Thermo Fisher Scientific 34080) and detected by photosensitization with ChemiDoc MP Imaging System (Bio-Rad). Primary antibodies are: Anti-BACH1 antiserum (A1-6 and 9D11 reported previously (18,27)), GAPDH (ab8245, Abcam), ACTB (GTX109639, GeneTeX), Phospho-p44/42 MAPK (9101, Cell Signaling Technology), p44/42 MAPK (9102, Cell Signaling Technology), Phospho-mTOR (pmTOR, SER2448) (2971, Cell Signaling Technology), mTOR (2972, Cell Signaling Technology)

### ChIP-sequence

The ChIP-DNA was prepared using anti-BACH1 antibody (A1-6) in HSC-2 or Ca9-22. Please refer to our previous reports for detailed protocols (18). The ChIP-DNA libraries were prepared from approximately 10 ng of ChIP-DNA or Input-DNA using the Ovation Ultralow DR Multiplex System (NuGEN Technologies Inc., CA, USA). The pooled libraries were sequenced as 51-base paired-end reads on an Illumina HiSeq 2500 (Illumina) across two lanes of rapid mode flow cells. The low-quality reads marked by Illumina sequencers in the fastq format data were filtered using CASAVA v1.8. Other lowquality reads were removed by Fastx toolkit v.0.0.14. The obtained reads were aligned to the hg19 human reference genome using bwa v 0.7.17. The peaks were called using MACS 2 v 2.1.2. The peaks existing within 100 bases in the different samples were judged as overlapping peaks.

### RNA-sequence

Five µg of the total RNA extracted by RNeasy plus mini kit (Qiagen) was treated with a RiboMinus Human/Mouse Transcriptome Isolation Kit (ThermoFisher scientific) to remove ribosomal RNA (rRNA). For fragmentation, 100 ng of the rRNA-depleted RNA was incubated at 95°C for 10 min and was purified by a Magnetic Beads Cleanup Module (Thermo Fisher Scientific).

Libraries were constructed using Ion Total RNA-seq Cartridges Beads Cleanup Module (Thermo Fisher Scientific). Templates were prepared using Ion PI Hi-Q Chef Kit (Thermo Fisher Scientific on Ion Chef (Thermo Fisher Scientific). Sequence runs in Ion PI Chip V3 (Thermo Fisher Scientific) were performed on Ion Proton (Thermo Fisher Scientific) using Ion PI Hi-Q Sequencing Kit (Thermo Fisher Scientific). Alignment of reads to reference hg19 and read count were performed using the RNASeqAnalysis plugin from the Ion torrent suite software. Count-based differential expression analysis was performed on EdgeR v3.30.3 after removal of low count lead genes using three biological replicates for each condition (less than 5 reads per gene in the sample and counts per million mapped reads (CPM) of 1 or less).

### Gene Ontology Analysis

GO analysis were performed using DAVID v6.8 on web site (https://david.ncifcrf.gov/summary.jsp).

### Generation of Tet-ON stable cell line

pRetroX-Tight-Pur (Clontech) was used as the tet-on vector. The human BACH1 sequence (hBACH1) was amplified by PCR using KOD-Plus-Neo (TOYOBO KOD-401) with primers having BamH1 (5ʹ-end) and EcoR1 (3ʹ-end) restriction enzyme cleavage sequences. The vector’s multi-cloning site and the end of the amplified hBACH1 were cleaved with BamH1 and EcoR1, and then ligated to construct pRetroX-Tight-Pur-hBACH1. Finally, we confirmed that there were no mistakes in the sequence of the insert hBACH1 using Applied Biosystems® 3500xL Genetic Analyzer (Thermo Fisher Scientific). The HSC-2 cell line stably expressing rtTA was generated by infecting cells with pMigR-rtTA-GFP/Bla-derived retrovirus and then drug-selecting with 20 µg/ml of Blasticidin S HCl (Thermo Fisher Scientific R210-01, rtTA-HSC-2). Then, tTA-HSC-2 was infected with pRetroX-Tight-Pur-hBACH1-derived retrovirus or pRetroX-Tight-Pur-derived virus as control. The stable cell lines were obtained by drug selection with 2 µg/ml puromycin dihydrochloride (Aldrich P9620-10ML, hBACH1-HSC-2, C-HSC-2).

### Administration of Tipifarnib, FeSOD and DFX

Tipifarnib (Selleck, R115777) was dissolved in DMSO. FeSOL was dissolved in 1 mM HCl (dissolved in distilled water). DFX was dissolved in DMSO All of these were diluted with RPMI-1640 and used.

### Cell death determination by flow cytometry

Cells were collected and pelleted and then, cells were suspended with 100 µl of each sample of 1×Annexin binding Buffer (BD Pharmingen™556454), and furthermore, 3 µl of APC-labeled Annexin V (BD Pharmingen™ 556454) was administered, and the cells were left to react for 5 minutes at room temperature in the dark. The cell suspension was then scale-up 5-fold by adding of 1× Annexin V binding buffer, and PI was added to a final concentration of 1 µg/mL and analyzed by flow cytometer (BD FACSVerse). Cells positive for both PI and Annexin V, or cells positive for only PI or Annexin V, were considered dead cells. The results were analyzed using the FlowJo™ v10 analysis software.

### Detection of labile iron ions by flow cytometry

Labile iron ions were detected using Mito-FerroGreen (Dojindo, M489). The cells were collected including medium used for culture and PBS used for washing, centrifuged, pelleted, lysed with 2ml HBSS (Hanks’ Balanced Salt Solution), washed thoroughly, centrifuged, and pelleted again. Mito-FerroGreen (1 mM in Ethanol) was added to the samples at a final concentration of 5 µM in HBSS, and the cells were incubated in a 37°C 5% COL incubator for 30 minutes. The cells were collected and stained for Annexin V and PI in the same way as the protocol used for cell death determination and then analyzed by flow cytometer (BD FACSVerse).

### Determination of Oxygen Consumption Rate

Oxygen consumption rate (OCR) was determined using Seahorse XF Analyzer (Agillent). Cells (6 x 10^5^) were seeded in 6 cm dish, treated with siRNA as described above after 24 hours, cultured for another 24 hours, treated with tripsine, and re-seeded onto microplates for the analyzer at 7.5 x 10^3^ cells/well in XF RPMI medium containing 1 mM pyruvate, 2 mM glutamine and 10 mM glucose. Where indicated, FeSO4 was added at 100 µM at the same time. Cells were cultured for another 1 hr and used for the measurement using XF Mito Stress Kit (Agillent), 0.1 µM oligomycin, 0.5 µM FCCP and 1.0 µM each of rotenone and antimycin A.

### Statistical analysis and plotting

Quantitative data were calculated as mean and standard deviation. For significance tests, one-way analysis of variance and Tukey method were used for multi-group comparisons; t-test was used for two-group comparisons; Student’s t-test or Welch’s t-test was used depending on the variance of the data. The significance level was set at 0.05, and the difference was considered statistically significant when the significance level was less than 0.05. GraphPad Prism8 was used for plotting and statistical analysis.

## Result

### Knockdown of BACH1 induces cell death in HNSCC cells

When BACH1 was knocked down by RNA interference in human HNSCC cell lines HSC-2 and Ca9-22, there was a significant increase in the number of floating cells and the percentage of cells adhering to the dish was decreased.The remaining adherent cells showed morphological changes including cytoplasmic swolling and enlarging. Many white vacuoles appeared in the cytoplasm and nucleus, and dark granular material was scattered or aggregated in the cytoplasm (Fig. 1A and B, see also Fig. 2D). The number of adherent cells decreased over time upon was knocked down (Fig. 1C). Flow cytometry analysis using Annexin V and propidium iodide (PI) revealed that the knockdown of BACH1 resulted in cell death (Fig. 1D).

**Figure 1.**
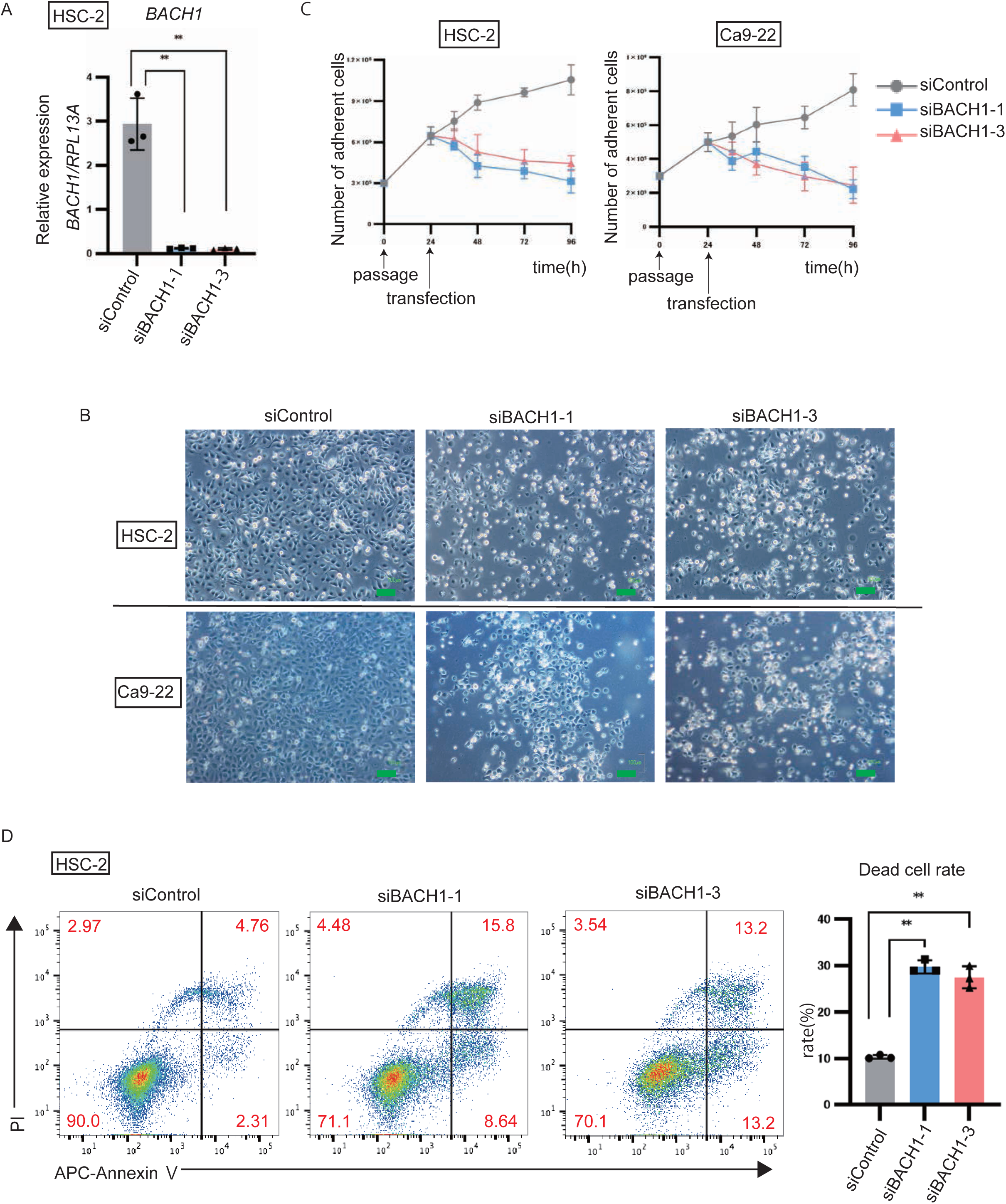
Head and neck cancer cells evade cell death depending on BACH1. (A) The expression levels of *BACH1* detected by RT-qPCR in HSC-2 cells with or without *BACH1* knockdown (two siRNAs). Each experiment was performed at least three times independently, and the column diagram shows the values for each experiment as dots, mean values as columns, and standard errors as bars. Statistical significance was tested by Student’s t-test, and p values are shown (** : p < 0.01). (B) Microscopic images of HSC2 and Ca9-22 cell morphology at 48 hours after *BACH1* knockdown. The scale represents 100 µm. (C) The cell numbers of HSC-2 and Ca9-22 with or without *BACH1* knockdown were manually measured over time to 12, 24, 48, and 72 hours after RNA interference, and the changes in cell proliferation are shown as a line graph. The experiment was performed three times independently, and the mean value and standard error are shown for each time point. (D) The percentage of dead cells in HSC2 and its control cells at 72 h after *BACH1* knockdown was measured by flow cytometry. A representative figure of gating is shown. Each experiment was performed at least twice independently, and three samples of each were prepared for each experiment. In the column diagram, values are shown as dots, mean values as columns, and standard errors as bars. Statistical significance was tested by Student’s t-test and p values are shown (** : p < 0.01). Allophycocyanin (APC) is shown on the X-axis and Propidium iodide (PI) on the Y-axis. Each sample was prepared in triplicate, and each experiment was performed at least twice independently.

**Figure 2.**
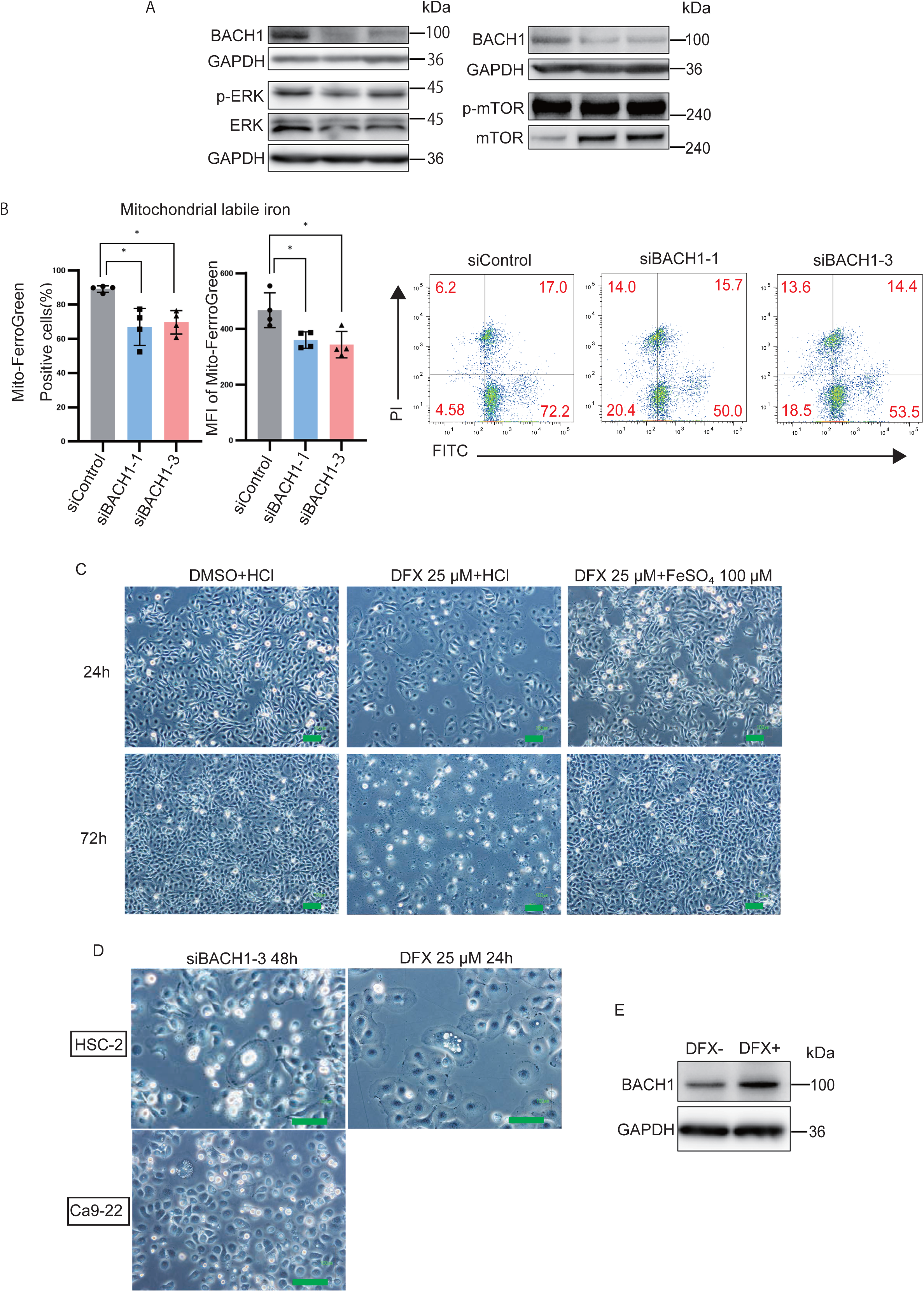
Intracellular labile iron is reduced with *BACH1* knockdown in HSC-2. (A) The expression levels of BACH1, GAPDH, ERK, mTOR, and the phosphorylation status of ERK and mTOR in HSC-2 cells were examined by Western blotting. Cells were subjected to total protein extraction 48 hours after transfection of siRNAs. In the left panel, 9D-11 was used as BACH1 antibody, and in the right panel, A1-6 was used as BACH1 antibody. Each experiment was performed at least three times independently. (B) Mitochondrial labile iron was detected by flow cytometry using Mito-FerroGreen fluorescein isothiocyanate (FITC) in HSC-2 cells with or without *BACH1* knockdown (24 h). The percentage of cells that become higher and the average fluorescence intensity (MFI) are shown. Each experiment was performed at least three times independently, and three or four of each sample were prepared for each experiment. The column plot shows values as dots, mean values as columns, and standard errors as bars. Statistical significance was tested by Student’s t-test and p-values are shown (* : p < 0.05). A representative gating diagram is shown on the right, with FITC on the X-axis and propidium iodide (PI) on the Y-axis. (C) Deferasirox (25 µM, dissolved in DMSO) and FeSOL (100 µM, dissolved in 1 mM HCl) were administered to HSC-2. DFX and FeSOL were administered simultaneously 24 h after cell passage. Microscopic images at 24 h and 72 h post-dose are shown. The scale represents 100 µm. (D) Microscopic images of HSC2 and Ca9-22 cells at 48 hours after *BACH1* knockdown and HSC-2 cells 24 hours after treatment of with 25 µM of DFX. The scale represents 100 µm. (E) Total protein extraction was performed at 24 h after the administration of 25 µM DFX to HSC-2, and the expression levels of BACH1 (A1-6 antibody) and GAPDH were examined by Western blotting. The experiments were performed independently at least three times.

### Head and neck cancers are addicted to labile iron maintained by BACH1

EGFR expression is one of the prognostic factors for salvage surgery for primary tumor recurrence in head and neck cancer (34). Since BACH1 is involved in ERK activation by repressing the expression of MAPK phosphatase (16), we examined the three signaling pathways downstream of EGFR, i.e., Ras/Raf/MEK/ERK, PI3K/AKT/mTOR, and JAK/STAT pathways. As judged by the phosphorylation of representative factors (ERK, mTOR, and STAT3), *BACH1* knockdown did not alter the status of ERK and mTOR in HSC-2 (Fig. 2A). The phosphorylation state of STAT3 was below the detection limit (data not shown). There was an increase in the amount of mTOR protein upon knockdown of *BACH1*. These results indicated that the cause of cell death upon *BACH1* knockdown could not be attributable to these signaling pathways.

Since BACH1 maintains the amount of labile iron which is not tightly bound to protein, in part by repressing the expression of ferritin genes (18–22), we next asked whether alterations in iron was responsible for the observed cell death upon BACH1 knockdown. We measured intracellular mitochondrial labile iron by flow cytometry using Mito-FerroGreen. The percentage of cells that were positive for Mito-FerroGreen was reduced in *BACH1* knockdown cells compared to control cells (Fig. 2B). The mean fluorescence intensity (MFI) was also decreased by knockdown of *BACH1* as well (Fig. 2B). These results showed that knockdown of BACH1 in HNSCC reduced intracellular labile iron.

The iron chelator Deferasirox (DFX) is known to cause cell death in several types of cancer cells (25–27, 32, 33). However, its effect on HNSCC cells has not been reported. Indeed, we found that DFX induced cell death in HSC-2 as well (Fig. 2C). And this cell death was suppressed by the addition of iron (II) sulfate (FeSOL), indicating the essentiality of iron in these cells. The morphological changes that occurred in the process of DFX were similar to those observed when *BACH1* was knocked down, including swelling of cytoplasm and cell enlargement, formation of vacuole-like structures in the cytoplasm and nucleus, and the presence of dark granular material scattered or aggregated in the cytoplasm (Fig. 2D). DFX increased the amount of BACH1 protein (Fig. 2F), suggesting the compensatory role of BACH1 in increasing cellular iron.

### Cell death upon BACH1 knockdown was rescued by iron

We next tested whether the cell death induced by knockdown of *BACH1* could be rescued by the addition of iron. When iron (II) sulfate was administered externally after knockdown of *BACH1* by RNA interference, cell death caused by knockdown of *BACH1* appeared suppressed (Fig. 3A). Quantification of the percentages of dead cells in HSC-2 and control cells with flow cytometry confirmed that the cell death was completely suppressed compared to the control (Fig. 3B). These results showed that cell death induced by knockdown of *BACH1* in HSC-2 is induced by cellular iron deficiency.

**Figure 3.**
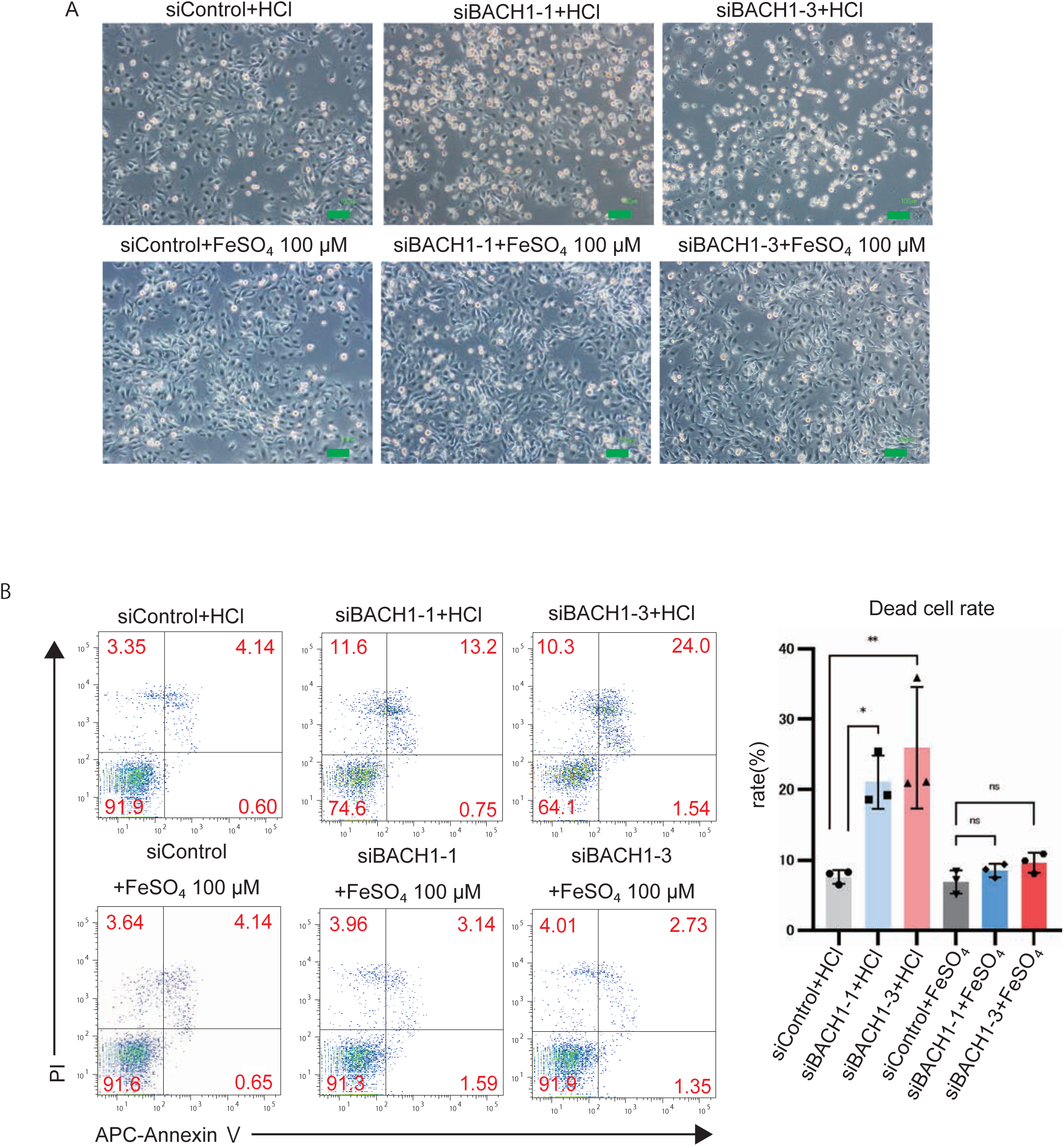
*BACH1* knockdown-induced cell death is completely rescued by iron administration. (A) Microscopic images of cell morphology in HSC2 and its control cells at 48 hours after *BACH1* knockdown are shown. FeSOL and HCl were administered 6 h after transfection. Scale indicates 100 µm. (B) The percentage of dead cells in HSC2 and its control cells at 48 hours after *BACH1* knockdown was detected by flow cytometry. A representative figure of gating is shown. Each experiment was performed at least twice independently, and three samples of each were prepared for each experiment. In the column diagram, values are shown as dots, mean values as columns, and standard errors as bars. Statistical significance was tested by Student’s t-test and p values are shown (* : p < 0.05, ** : p < 0.01). A representative gating diagram is shown on the left side, with APC on the X-axis and PI on the Y-axis.

### BACH1 directly represses ferritin gene expression in head and neck cancer

We then extracted RNA from *BACH1* knockdown HSC-2 and checked the expression of genes related to iron metabolism by real-time qPCR. Among those genes, the expression of both ferritin genes (*FTL* and *FTLH1)* was both increased in *BACH1* knockdown HSC-2 (Fig. 4A). The expression of transferrin receptor gene (*TFRC*) was increased by *BACH1* knockdown (Fig. 4A), suggesting that iron uptake into the cells was enhanced to compensate for the iron deficiency. Ferroportin (*SLC40A1*), which exports iron out of cells, was below the detection limit (data not show). To validate its role in HSC-2 cells, we generated cells with inducible expression of BACH1 (Fig. 4B). Upon BACH1 induction, *FTL* was significantly downregulated, while *FTH1* did not show significantl alteration (Fig. 4 C).

**Figure 4.**
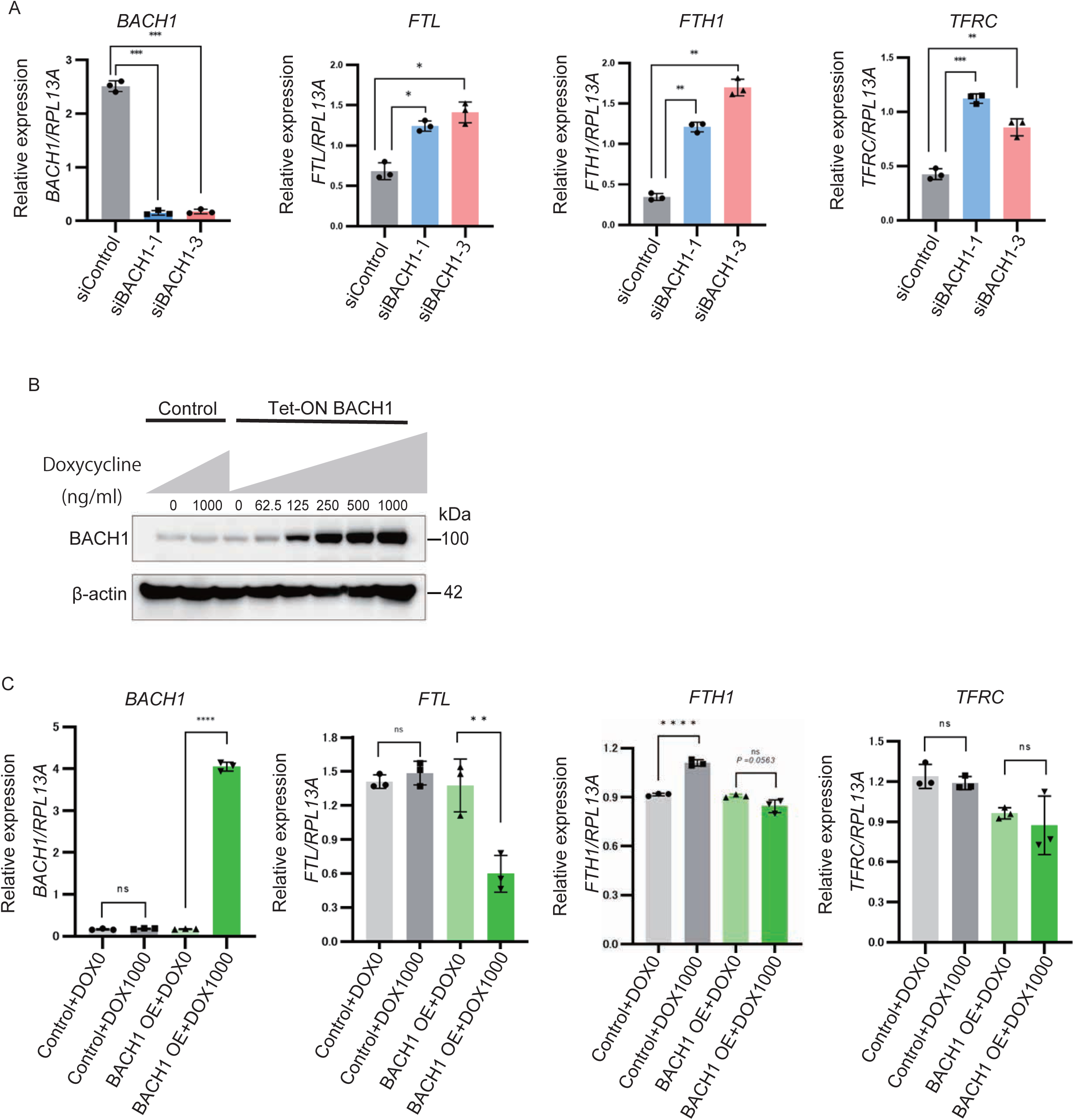
BACH1 represses ferritin gene expression in HNSCC. (A) The expression levels of *BACH1*, *FTL*, *FTH1*, and *TFRC* detected by RT-qPCR in HSC-2 cells with or without *BACH1* knockdown. Each experiment was performed at least three times independently, and the column diagram shows the values for each experiment as dots, mean values as columns, and standard errors as bars. Statistical significance was tested by Student’s t-test, and p values are shown (* : p < 0.05, ** : p < 0.01, *** : p < 0.001). (B) A retroviral vector incorporating the coding region of *BACH1* downstream of a promoter inducible with Doxycycline was generated. Using HSC-2, cells infected with this vector (TET-on BACH1 HSC2) and cells containing the control vector (Control HSC-2) were prepared and cultured at varying doses of Doxycycline. Total protein extraction of the samples added at each Doxycycline concentration was performed, and BACH1 (using A1-6 as antibody) and GAPDH were detected by Western blotting. The unit for each Doxycycline value is ng/ml. (C) Expression levels of *BACH1*, *FTL*, *FTH1*, and *TFRC* detected by RT-qPCR in Tet-on BACH1 HSC-2 and control HSC-2 cells without and with 1000 ng/ml of Doxycycline, respectively. Each experiment was performed at least three times independently, and the column diagram shows the values for each experiment as dots, mean values as columns, and standard errors as bars. Statistical significance was tested by Student’s t-test and P values are shown (** : p < 0.01, **** : P < 0.0001).

To capture global view of BACH1 function in HNSCC cells, we performed chromatin immunoprecipitation sequencing (ChIP-seq) of anti-BACH1 antibodies using HSC-2 and Ca9-22 cells. We also analyzed gene expression in HSC-2 cells with or without *BACH1* knockdown using RNA-sequence (RNA-seq) analysis. The binding sites of BACH1 in HNSCC cells were less than those in pancreatic cancer cells we reported previously (Fig. 5A)(12), which may reflect the amounts of BACH1 proteins in these cells. The shared peaks of HNSCC and pancreatic cancer cells were mapped in the vicinity of 403 genes and included genes for ferritin (*FTL*, *FTH1*) (Fig. 5A and B). These results suggested that BACH1 directly repressed the transcription of both *FTL* and *FTH1* genes in HNSCC to increase labile iron. Gene ontology (GO) analysis of these genes showed enrichment of genes related to cell contacts and motility (Fig. 5C), suggestive of its involvement in metastasis as well. When combined with the results of RNA-seq analysis, we found that, while the principal function of BACH1 is as a repressor of transcription, more numbers of genes were down regulated upon BACH1 knockdown in these cells (Fig. 5D). Multiple genes involved in iron metabolism were also reduced in their expression (Fig. 5D). GO enrichment analysis of the clusters of the up-regulated and down-regulated genes showed several characteristic changes which are consistent with the observed effects of BACH1 knockdown (Fig. 5E). For example, genes in “positive regulators of apoptosis” and “negative regulators of cell proliferation” were increased in their expression, which may be related to the process of cell death observed in this study. The reduced expression of genes related to “oxidation-reduction process”, many of which are iron-dependent enzymes, may reflect compensatory responses to the reduction of labile iron by the knockdown of BACH1. The decreased expression of genes in “positive regulation of cell migration” is consistent with previous report that knockdown of BACH1 in pancreatic cancer cells results in decreased migration and metastasis (12). Decrease in the expression of genes related to cholesterol synthesis was also found. As iron induces cholesterol synthesis (35, 36), which is important for cell proliferation, the decrease in labile iron upon knockdown of *BACH1* may reduce cholesterol synthesis. To extract genes that are likely to be directly regulated by BACH1 in HNSCC cells, we merged the ChIP-seq results of BACH1 with the transcripts that were up-and down-regulated by more than 1.5-fold when BACH1 knockdown in HSC-2 cells. We found 199 repressed genes and 157 activated genes (Fig. 5F). These genes are involved in morphogenesis, immune response, PI-3 kinase signaling, and cell polarity (Fig. 5G), which may contribute to cancer cell properties.

**Figure 5.**
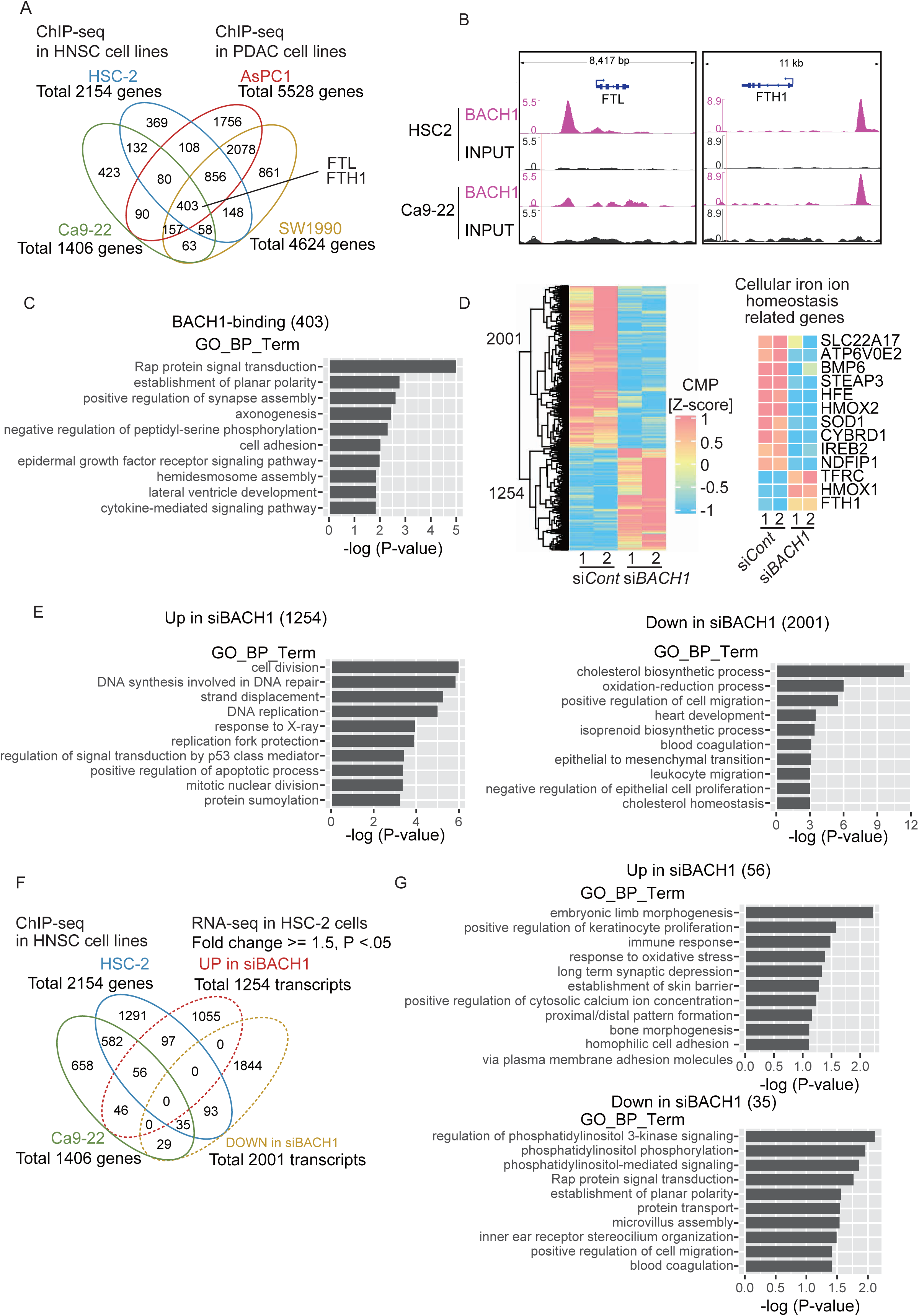
Direct target genes of BACH1 in HNSCC. A) Venn diagram merging the results of ChIP-seq with anti-BACH1 antibody in HNSCC cell lines HSC-2 and Ca9-22 with the previously reported results of ChIP-seq with anti-BACH1 antibody in pancreatic cancer cell lines AsPC1 and SW19901 (9). B) Visualization of the accumulation of BACH1 bound around the ferritin genes in HSC-2 and Ca9-22 using the Genome Viewer (IGV). C) Top 10 results of GO_BP analysis of 403 genes with nearby binding of BACH1. D) Heatmap of genes that were significantly (p < 0.05) and 1.5-fold or more differentially expressed in BACH1 knockdown HSC-2 compared to the control, with 1254 up-regulated and 2001 were down-regulated transcripts in BACH1 knockdown HSC-2. These included genes related to iron ion homeostasis in cells, as shown on the right. E) The results of GO analysis of genes whose expression was increased (left panel) or decreased (right panel) with BACH1 knockdown in HSC-2. F) Set analysis of ChIP-seq results of BACH1 and RNA-seq results in HSC-2. G) GO analysis of candidate direct target genes from F.

### BACH1 maintains labile iron for evading cell death and mitochondrial failure

To understand the types of cell death induced upon BACH1 knockdown, we treated control and knockdown cells with inhibitors of known cell death regulators (Fig. 6A). The inhibitors of apoptosis (Z-VAD) and necroptosis (necrostatin-1) showed recoveries of cell proliferation, and the combination of the two inhibitors fully rescued the effect of BACH1 knockdown. In contrast, ferroptosis inhibitor ferrostatin-1 did not show any effects. Therefore, the cells appeared to die due to a combined execution of apoptosis and necroptosis.

**Figure 6.**
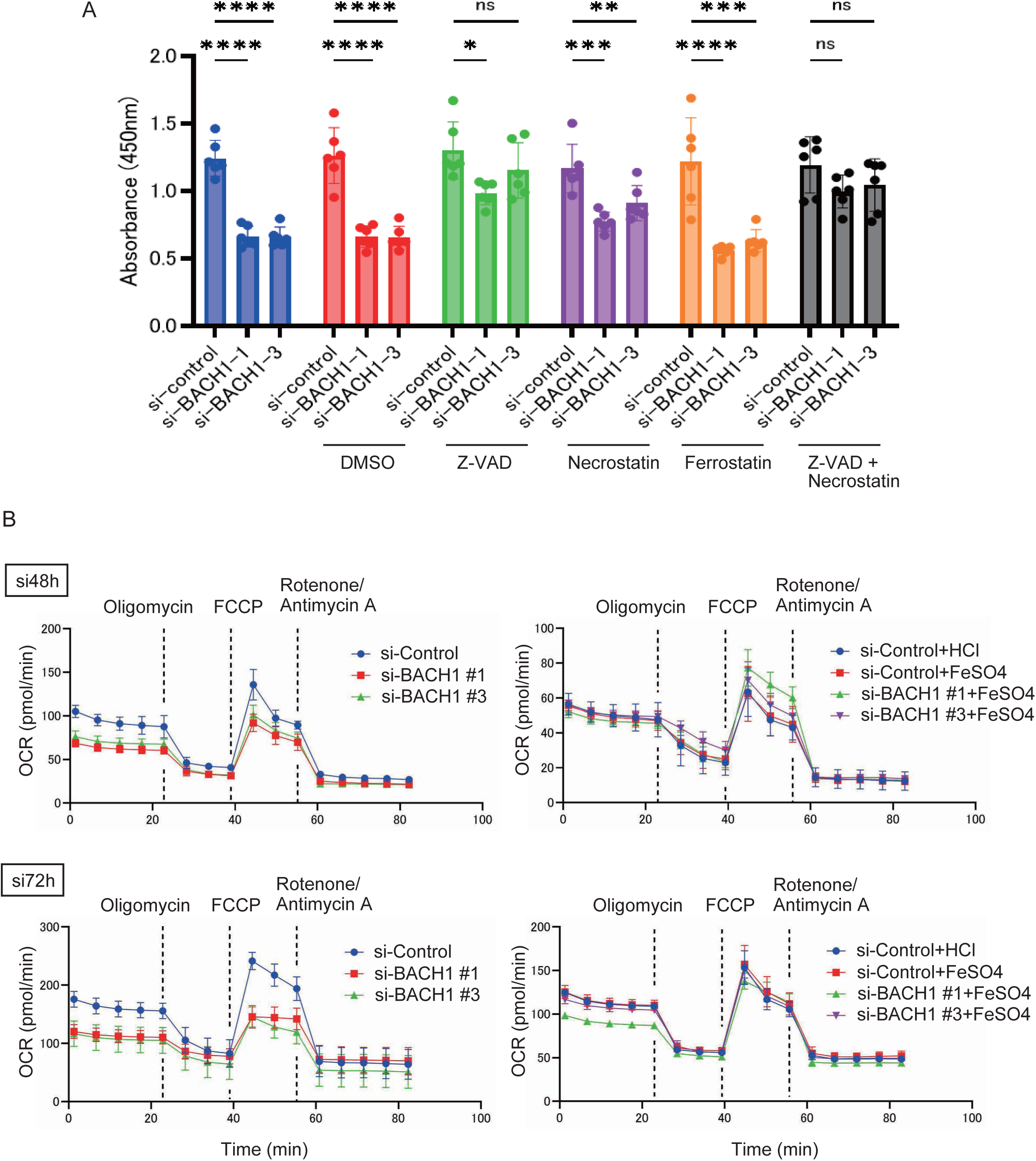
Mitochondrial defects in BACH1 knockdown. (A) HSC-2 treated with BACH1 siRNA were treated with indicated inhibitors of cell death and relative numbers of surviving cells were determined. ANOVA was used for the statistical test (* : P < 0.05, ** : P < 0.01, *** : P < 0.001). (B) Oxygen consumption rates of HSC-2 treated with control or two different BACH1 siRNAs were determined after 48 or 72 hours (left and right, respectively) after the introduction of siRNAs without or with FeSO_4_ (upper and lower).

Considering that these types of cell death often involve mitochondrial failure (37) and that iron is critical for healthy mitochondria (38), we examined the electron transfer chain activity as a read out of mitochondria activities (Fig. 6B). BACH1 knockdown reduced both basal level of respiration as well as maximal respiratory capacity. Importantly, these defects were completely rescued with the addition of iron to the culture medium. When taken together, these results indicated that the labile iron deficiency in BACH1 knockdown cells culminates in cell death of metabolic disorganization involving mitochondrial failures.

### Knockdown of BACH1 may induce sensitivity to Tipifarnib

Since the observed response of HNSCC cells to BACH1 knockdown involved alterations in iron metabolism and mitochondrial functions, the manipulation of BACH1 may synergize with therapeutic drugs targeting other mechanisms. Indeed, the combinations of DFX with cisplatin in esophageal cancer cell lines (32) and with gemcitabine in pancreatic cancer cell lines (33) are known to cause synergistic growth inhibition. However, there has been no report on the combination of DFX with Tipifarnib. We combined Tipifarnib with BACH1 knockdown. While HSC-2 do not carry activating mutations in *HRAS* (39), preclinical studies have shown that increased concentration of tipifarnib has weak antitumor effect on HNSCC cells (40). When HSC-2 cells were treated with different concentrations of tipifarnib, weak cytotoxic effect was observed at higher concentrations (Fig. 7A). After treating the control and *BACH1* knockdown HSC-2 cells with 10 nM Tipifarnib, which did not show any cytotoxic effect on the cells (Fig. 7B), we quantified by flow cytometry the percentages of dead cells (Fig. 7C). While Tipifarnib alone did not increase cell death, the combination with BACH1 knockdown increased cell death more than the knockdown alone. We concluded that *BACH1* knock down enhances the effect of Tipifarnib in HSC-2 cells.

**Figure 7.**
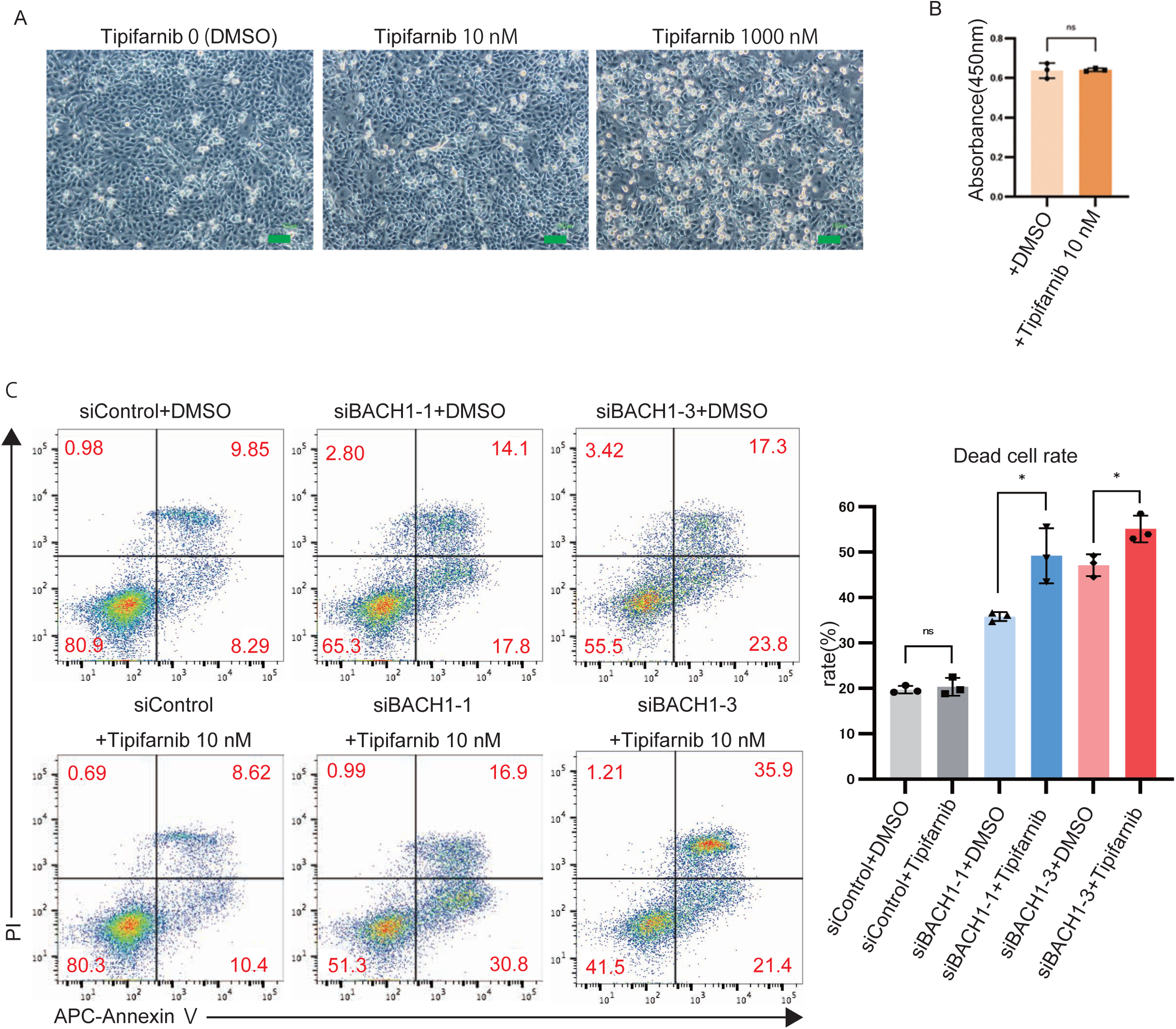
Knockdown of *BACH1* increases tipifarnib sensitivity. (A) Microscopic images of HSC-2 treated with Tipifarnib. The cells were treated with DMSO or Tipifarnib at 24 hours after cell passage and incubated for another 48 hours. Tipifarnib is dissolved in DMSO, and the amount of DMSO corresponds to that of 1000 nM Tipifarnib. Scale indicates 100 µm. (B) The cell activity of HSC-2 treated with DMSO or 10 nM Tipifarnib was evaluated by the WST method. HSC-2 cells were seeded on 96well plates (2000/well) and treated with DMSO or Tipifarnib after 24 hours, followed by 48 hours of culture, and relative cell numbers were determined. Nine samples of each were prepared, and the same experiment was performed three times independently. The mean value of each time was used as a dot, the mean value as a column, and the standard error as a bar. Statistical significance was tested by Student’s t-test. (C) DMSO or 10 nM Tipifarnib was administered 24 hours after *BACH1* knockdown, and the percentage of dead cells in HSC2 and their control cells was detected by flow cytometry at 48 hours after Tipifarnib administration. Each experiment was performed at least three times independently, and three samples of each were prepared for each experiment. In the column diagram, values are shown as dots, mean values as columns, and standard errors as bars. Statistical significance was tested by Student’s t-test and p values are shown (* : p < 0.05). A representative figure of gating is shown on the left, with APC on the X-axis and PI on the Y-axis.

## Discussion

In this study, we found that HNSCC cells are dependent on BACH1 in maintaining the homeostasis of intracellular labile iron for their survival. On knocking down BACH1 or treating with iron chelators, these cells underwent cell death involving both apoptosis and necroptosis. Because the observed cell death was efficiently suppressed by the supplementation of iron, the primary cause of the cell death is iron deficiency. Importantly, we showed that this sensitivity of HNSCC cells to iron metabolism could be exploited as a combination with RAS targeting drug tipifarnib.

One of the unexpected findings was that BACH1 knockdown caused iron deficiency in HNSCC cells even when cultured under iron-rich culture medium. BACH1 has been deleted or knocked down in various types of cells and in mice, but, under the normal experimental conditions, no report described iron deficiency or downstream changes due to iron deficiency. BACH1 knockout mice show defects in red blood cell formation only after being fed with iron deficient diet (41). Why do HNSCC cells show the clear dependence on BACH1 to maintain iron? Interestingly, ferritin was directly repressed by BACH1 in pancreatic cancer cells just like in HNSCC cells. Hence, increased iron storage by ferritin in response to BACH1 knockdown may be the same for these two types of cancers. Rather, HNSCC cells may require more labile iron than other types of cells. For example, they may be heavily dependent on electron transfer chain for ATP synthesis. Alternatively, iron flux in these cells may be slower than other types of cells. If iron uptake is slow, capturing of labile iron by excess ferritin is expected to lead to a rapid reduction of labile iron. Recycling and supply of labile iron via ferritinophagy may be slow in HNSCC cells. In this case, even if total cellular iron is high, usability of iron could become low, like non-circulating money. It is noteworthy that the use of iron chelators to deplete iron and the use of iron-deficient diets have already been reported to be useful in cancer treatment in clinical practice (25, 42–45). Therefore, the control of intracellular labile iron may pose a novel therapeutic strategy in cancer treatment including HNSCC. Whether BACH1-dependency in iron metabolism is specific to a subset of cancers like HNSCC is a subject for further investigation.

How the labile iron deficiency would lead to cell death upon BACH1 knockdown in HNSCC cells? Ferroptosis, a form of cell death induced by reactive labile iron, is characterized by iron-dependent lipid peroxidation and is inhibited by iron chelators (45). These features are not consistent with the cell death observed in this study. Iron chelators such as deferasirox and deferoxamine can also induce cell death under certain conditions (26, 44, 46). It has been reported that iron chelators induced ROS production in progenitor cells of acute myeloid leukemia, resulting in cell death (27). Others have reported that iron deficiency inhibits DNA synthesis by reducing the activity of ribonucleotide reductase which is involved in deoxyribonucleic acid synthesis (47, 48). Iron deficiency also suppress the citric acid cycle and electron transfer chain systems in mitochondria, leading to reduced cellular energy production, because many of the proteins and enzymes involved are iron-dependent (49). Iron deficiency in mitochondria was shown to promote mitophagy, a subtype of autophagy, resulting in mitochondrial dysfunction and cell death (44). Because of the plethora of biochemical reactions that depend on iron, it is difficult to narrow down critical iron-dependent proteins or reactions in the present study. Nonetheless, the clear reduction of electron transfer chain activity upon BACH1 knockdown, together its rescue with iron supplementation, strongly points to mitochondria as the downstream function which is strictly dependent on BACH1 and hence labile iron in HNSCC cells.

BACH1 protein was up-regulated in HSC-2 cells when treated with DFX. This is consistent with a feedback mechanism that prevents excessive depletion of labile iron; during intracellular labile iron deprivation, the expression of BACH1 is increased, thereby suppressing ferritin genes expression and consequently preserving labile iron. Thus, it is speculated that BACH1 is closely involved in iron homeostasis in HNSCC cells. Heme-dependent ubiquitination of BACH1 (50) can mediate such a regulation. More directly, iron itself may regulate BACH1, which is an important avenue of future research.

The integrated analysis of BACH1 ChIP-seq and RNA-seq allowed us to grasp global view of the BACH-dependent gene program in HNSCC cells. We found reduced the expression of genes involved in oxidation-reduction process, which includes many of the iron-dependent enzymes. This may reflect an indirect, adaptative response to the labile iron deficiency caused by BACH1 knockdown. Genes related to cholesterol synthesis was also decreased. Since cholesterol synthesis is important component of cell membranes, a reduction in cholesterol may lead to cellular fragility and/or reduced mitochondrial function. In addition, Genes in PI3-kinase and Rap signaling cascades are down regulated. These changes in gene expression may also contribute to the observed cell death of *BACH1* knockdown cells.

The limitation of tipifarnib is that it has limited efficacy on tumors with fewer alle frequency of *HRAS* mutations and those without *HRAS* mutations (9). In xenograft studies of melanoma cell lines, it has been reported that high doses of tipifarnib can inhibit tumor growth even in the absence of *HRAS* mutations (40). However, increasing the drug concentration also increases adverse events in clinical practice; there is a need to find a way to provide a high anti-tumor effect within the range of practical dosage. In this study, we have shown that knockdown of *BACH1* can confer Tipifarnib sensitivity to HSC-2 cells lacking *HRAS* mutation. Besides the genes related to iron metabolism, the knockdown of BACH1 caused decreased expression of downstream signaling cascades of RAS, including RAP and PI3-kinase pathways. These alterations may also contribute to the enhancement of tipifarnib effect. We previously reported that *Bach1* is required for transformation and tumorigenesis of mouse embryonic fibroblasts (MEFs) with mutant *Hras* and tumor formation in chemical carcinogenesis model of oral cancer (16). These observations suggest that BACH1 inhibition will potentiate the effect of Tipifarnib on HNSCC.

In conclusion, we found that BACH1 plays an important role in the homeostasis of intracellular labile iron in HNSCC cells which poses a new metabolic vulnerability of these cells. It remains to be elucidated whether other types of cancer cells are also responsive to BACH1 inhibition and labile iron perturbation. Together with previous reports (12, 16, 18–23), it is now clear that BACH1 promotes cancer cell progression by regulating multiple pathways including iron metabolism, mitochondrial functions, glycolysis and EMT.

## Authors’ Contributions

Conceptualization: M. Rokugo, K. Nakamura, M. Matsumoto, N. Long, K. Igarashi

Investigation and data curation: M. Rokugo, K. Nakamura, M. Matsumoto, N. Long, H. Nishizawa, A. Endo, R. Shima, R. Funayama

Methodology: M. Rokugo, K. Nakamura, M. Matsumoto, K. Nakayama, T. Abe

Formal analysis: M. Rokugo, K. Nakamura, M. Matsumoto, N Long

Resources, funding acquisition: M. Matsumoto, K. Igarashi

Supervision: M. Matsumoto, A. Nakanome, A. Okoshi, T. Ogawa, Y. Katori, K. Igarashi

Writing – Original Draft Preparation: M. Rokugo, M. Matsumoto

Writing – Review & Editing: M. Matsumoto, K. Igarashi

## Acknowledgements

We thank members of the Departments of Biochemistry, Tohoku University Graduate School of Medicine for discussions and support, and the Biomedical Research Core of Tohoku University Graduate School of Medicine for technical support.

## Funding information

This study has been supported by Grants-in-Aid for Scientific Research from the Japan Society for the Promotion of Science (22H00443, 20KK0176 and 18H04021 to KI, 19K07680 and 16K07108 to MM, and 20K16296 and 19K23738 to HN), Grant-in-Aid for Joint Research by Young Researchers from Tohoku University Graduate School of Medicine (to H.N.), grants from the Japan Agency for Medical Research and Development (AMED, 24zf0127001h0004, 24fk0108655h0003 and JP24fk0210114 to T.A), Gonryo Medical Foundation (to H.N.), Takeda Science Foundation (to H.N.), Casio Science Promotion Foundation and Research Grant in the Natural Sciences from the Mitsubishi Foundation (to K.I.).

## Conflicts of Interest

No potential conflicts of interest were disclosed.

## Notes

### Competing Interest Statement

The authors have declared no competing interest.

